# A conserved cysteine–histidine–glutamate metal site identifies DUF501 (Rv1025), an essential uncharacterised protein family of *Mycobacterium tuberculosis*, as a candidate metalloenzyme and drug target

**DOI:** 10.64898/2026.08.31.746906

**Authors:** Christophe Guyeux

**Affiliations:** Femto-ST Institute, UMR 6174 CNRS, Université Marie et Louis Pasteur, Besançon, France

**Keywords:** *Mycobacterium tuberculosis*, drug target, DUF501, metalloenzyme, AlphaFold, protein family characterisation, dark proteome

## Abstract

A substantial fraction of the *Mycobacterium tuberculosis* proteome remains functionally uncharacterised. Rv1025, a 155-residue protein carrying the domain of unknown function DUF501 (Pfam PF04417), is essential by transposon mutagenesis and vulnerable by CRISPR interference, an attractive but neglected drug target, yet has never been functionally described. The family (4,370 proteins, no Gene Ontology term, no solved structure) is uncharacterised across all organisms and essential in three Actinobacterial genera. A Foldseek search of the AlphaFold model against complete structural databases finds no significant homolog, indicating a novel fold. The operon *eno*–*divIC* –*Rv1025* –*ppx2* is conserved across the Actinobacteria phylum, yet AlphaFold-Multimer finds no direct complex between Rv1025 and its neighbour DivIC. Instead, conservation across 8,700 homologous sequences reveals a near-invariant Cys113–His115–Glu59 cluster forming a pocket. Holo AlphaFold3 predictions with Zn, Fe and Mn confidently place a divalent metal on this triad at 2.25–2.47 Å; mutating the triad relocates the metal, and an independent backbone-geometry predictor recovers the same site, confirming specificity. The triad is universal across the family: present in all 1,472 near-complete bacterial sequences of the Pfam alignment, with no non-conservative substitution among the 2,228 sequences examined, a defining feature of bacterial DUF501 rather than a mycobacterial peculiarity. We propose that DUF501 is a metal-binding protein and candidate metalloenzyme, the first functional hypothesis for this family, whose conserved, essential metal pocket is a promising drug target. As the predictions build on a conservation-defined site within a fully computational study, they are supportive rather than proof of metal occupancy and warrant experimental validation.

## 1 Introduction

Tuberculosis remains a leading cause of death from a single infectious agent, and the length of curative treatment together with the spread of drug resistance sustains a pressing need for antibiotics acting on new targets. Twenty-five years after the *Mycobacterium tuberculosis* H37Rv genome was completed (Cole et al., 1998), a large part of its proteome is still annotated as “conserved hypothetical” or carries only a domain of unknown function (DUF). Some of these proteins are not dispensable curiosities: genome-wide transposon mutagenesis (DeJesus et al., 2017) and CRISPR-interference vulnerability screens (Bosch et al., 2021) show that a subset are both essential for growth and highly sensitive to partial knockdown, precisely the profile sought for a drug target. Because they are uncharacterised, they are systematically overlooked.

Rv1025 is a 155-residue protein of this class. It carries a single domain, DUF501 (Pfam PF04417, InterPro IPR007511, orthologous group COG1507, KEGG orthology K09009), and no enzymatic or functional annotation. It sits in a four-gene operon, *eno* (Rv1023, enolase)– *divIC* (Rv1024, a cell-division protein of the FtsB/DivIC family)–*Rv1025* –*ppx2* (Rv1026, an exopolyphosphatase (Chuang et al., 2015; Choi et al., 2012; Tiwari et al., 2019)). The heterogeneity of these neighbours (glycolysis, division, polyphosphate metabolism) has so far prevented any pathway assignment. Yet Rv1025 is essential (DeJesus et al., 2017), strongly vulnerable to knockdown, mass-spectrometry detected, near-invariant across clinical isolates, and predicted to be very well folded, all of which single it out as a tractable candidate target that has never been studied for its own sake.

Here we combine population-scale conservation, structural search, genomic synteny across the Actinobacteria, and AlphaFold3 complex and holo modelling to propose the first functional hypothesis for DUF501. We find that DUF501 adopts a fold with no known structural relative, that its operon is conserved across the entire phylum, that Rv1025 does not form a stable binary complex with DivIC despite this synteny, and that a near-invariant Cys–His–Glu triad forms a metal-binding site, supported by holo AlphaFold3, that we interpret as a candidate, possibly catalytic centre.

## 2 Materials and Methods

### Sequence, annotation and conservation

The Rv1025 protein sequence (UniProt P96375, 155 aa) and its operon neighbours were taken from the H37Rv reference (NC_000962.3) (Cole et al., 1998). Family-level statistics (member and taxon counts, Gene Ontology terms, associated PDB structures) were retrieved from InterPro/Pfam (Paysan-Lafosse et al., 2023) and the orthology from eggNOG (Huerta-Cepas et al., 2019). Essentiality and vulnerability were taken from published transposon (DeJesus et al., 2017) and CRISPRi (Bosch et al., 2021) screens; cross-genus essentiality was obtained from the *Corynebacterium diphtheriae* essential-genome study (Goodall et al., 2023). Per-residue conservation was computed from the deep multiple-sequence alignment generated by AlphaFold3 for Rv1025 (8,700 sequences); after removing insertion states, each query position was scored by the fraction of non-gap sequences matching the query residue.

### Structure model and structural search

The AlphaFold model of Rv1025 (AlphaFold DB v6, AF-P96375-F1, mean pLDDT 95.3) (Jumper et al., 2021; Varadi et al., 2022) was searched with Foldseek (van Kempen et al., 2024) against the complete PDB100, AlphaFold/Swiss-Prot and CATH50 databases (mode 3Di+AA). A complementary profile–profile (HMM–HMM) search was run with HHpred (Zimmermann et al., 2018) against PDB_mmCIF70, Pfam-A and SCOPe70.

### Synteny across Actinobacteria

The four operon proteins were queried by `tblastn` (Camacho et al., 2009) (e-value ≤ 10^−5^) against 63 reference genomes: 48 non-tuberculous *Mycobacterium* species, *M. canettii* and *M. bovis* as controls, ten non-*Mycobacterium* Actinobacteria spanning Corynebacteriales, Streptomycetales, Bifidobacteriales, Propionibacteriales and Micrococcales, and *Escherichia coli* and *Bacillus subtilis* as out-of-phylum negatives. For each genome the best hit per query was retained; synteny was assessed by co-localisation and gene order around the DUF501 anchor. To test whether the operon’s synteny exceeds the genome-wide baseline, we drew 90 random four-gene, co-directional H37Rv blocks and scored each, by the same `tblastn` procedure, across the eleven distant (non-*Mycobacterium*) Actinobacteria. To separate arrangement conservation from gene conservation, scoring was conditioned on orthology: for each block we counted the genomes in which all four orthologs were found (present) and, among those, the genomes in which they were syntenic (single contig, span ≤ 8 kb, consistent order); the per-block statistic is the fraction syntenic given present, and the operon was placed in the resulting null distribution (empirical *p* from the fraction of random blocks at least as syntenic). A schematic of the block along the Actinobacterial taxonomy was drawn with Matplotlib (Supplementary Figure S3).

### Complex and holo modelling

AlphaFold3 (Abramson et al., 2024) was run through the AlphaFold Server with a fixed model seed, returning five models per job. Putative interactions were tested as heterodimers (Rv1025 with DivIC; specificity controls Rv1025 with Enolase and with Ppx2; positive control DivIC with the divisome protein FtsQ, Rv2151c) and read out by interface pTM (ipTM) and the minimum inter-chain predicted aligned error, using the five models per job. The metal site was tested by holo predictions of Rv1025 with one Zn, Fe or Mn ion; coordination was measured on the top-ranked model as the distances between the ion and the candidate donor atoms, with the per-ion pLDDT as a confidence estimate. As a negative control, the three ligands were mutated to alanine (C113A/H115A/E59A) and the holo prediction repeated. As a robustness check of the DivIC result, five further divisome components (FtsZ, FtsW, PbpB/FtsI, FtsK, SepF) were screened as heterodimers with Rv1025 using Boltz-2 (Passaro et al., 2025) run locally on CPU, reusing the Rv1025 multiple-sequence alignment generated by AlphaFold3 and the DivIC–FtsQ pair as an in-panel positive control; because Boltz-2 single-model CPU inference is a distinct, noisier predictor from the AlphaFold Server, these results are reported separately and are not pooled with the AlphaFold3 figures. A direct Rv1025 homodimer, with and without the two holo Fe ions, was likewise tested by AlphaFold3 to ask whether a second subunit could complete the metal coordination sphere in trans. Structure parsing used Biopython (Cock et al., 2009); molecular figures were rendered with PyMOL.

### Family-wide conservation of the triad

The full Pfam alignment of PF04417 (2,228 sequences, 471 columns) was retrieved from the InterPro API and used as provided, without realignment. Because Rv1025 is itself a member (P96375/16–137), the alignment columns corresponding to Glu59, Cys113 and His115 were located by walking its aligned row, and each sequence was then read at those three columns. Sequences were called near-complete when their aligned length reached 90% of the median length of triad-bearing members, which separates genuine substitutions from gaps in fragments; conservative substitution was defined as retention of the coordinating chemistry (Asp for Glu). Taxonomic lineages were obtained from the UniProt REST API for every accession, and identity to Rv1025 was computed over columns where both sequences are ungapped.

### Coevolution analysis

Residue coevolution was computed on the same 8,700-sequence alignment by mean-field direct coupling analysis (Morcos et al., 2011): sequences were reweighted at 80% identity, frequencies regularised with a 0.5 pseudocount, and couplings obtained by inverting the covariance matrix, converted to Frobenius norms in the zero-sum gauge and corrected for background with the average product correction (Dunn et al., 2008). Columns with more than 50% gaps and pairs separated by fewer than five residues were excluded. Predicted pairs were compared with the apo model using C*β*–C*β* distances, contacts defined at 8 Å; pairs beyond 15 Å were treated as not explained by the monomer. Relative solvent accessibility used Shrake–Rupley areas normalised to Tien et al. (2013) maxima. Enrichment of couplings between the two pockets was tested against a distance-matched null of 10,000 resamplings, since coupling strength decays with distance.

### Orthogonal metal-site prediction and druggability

To assess the metal site independently of the conservation signal and of AlphaFold’s learned metal placement, the apo model was analysed with BioMetAll (Sánchez-Aparicio et al., 2021), which identifies metal-binding sites from backbone preorganisation alone, and with AlphaFill (Hekkelman et al., 2023), which transplants ligands and ions from homologous holo structures. Pocket detection and druggability scoring of the apo model used fpocket (Le Guilloux et al., 2009); each detected pocket was mapped to the metal site by the minimum distance between its alpha-sphere centres and the Cys113/His115/Glu59 donor atoms.

### Metal-site accessibility

Because fpocket’s hydrophobicity-weighted scoring function is not designed for polar metal sites, accessibility of the coordinated ion was instead measured geometrically, independently of that scoring function. On each of the five holo models per ion (Zn, Fe, Mn; water and any other non-protein atom excluded throughout), 4,000 directions were cast from the ion at 2.1 Å; a direction was scored *open* if no protein atom lay within 2.4 Å of the probe (repeated at 2.6 and 2.8 Å thresholds as a robustness check) and *solvent-facing* if it additionally reached 10 Å from the ion without passing within 1.9 Å of any atom. The same procedure was applied, with the identical thresholds, to four calibration structures: three validated catalytic zinc sites (human carbonic anhydrase II, PDB 1CA2 (Eriksson et al., 1988); thermolysin, PDB 8TLN (Holland et al., 1992); the bacterial deacetylase LpxC, PDB 4MDT (Clayton et al., 2013)) and one structural, non-catalytic zinc site (the Zif268 zinc-finger domain, PDB 1ZNF (Lee et al., 1989)). To separate genuine accessibility from a possible bias of AlphaFold models toward looser packing than crystal structures, the same measurement was repeated on the AlphaFold DB models of the two calibration proteins with a crystal-derived active site (carbonic anhydrase II, thermolysin), the metal transplanted onto the model by Kabsch superposition of paired C*α* atoms after resolving any numbering offset between the crystal and UniProt sequences.

The bioinformatic analyses, database queries and literature cross-referencing underlying this study were carried out within an AI-augmented bioinformatics working environment, Claude Code (Anthropic), operating the Claude Sonnet 5 language model as an interactive analysis agent under continuous human direction. Every script, statistical test and numerical claim produced within this environment was independently traced back to its underlying raw data, primary publication or database record before being reported here, following the verification discipline described throughout this section; the corresponding author designed the study, specified and reviewed every analysis, and takes full responsibility for the accuracy of its content.

## 3 Results

### 3.1 Rv1025 and the DUF501 family are essential yet uncharacterised across Actinobacteria

Rv1025 is a genuine, well-supported gene product: it is essential by transposon mutagenesis (DeJesus et al., 2017), strongly vulnerable to CRISPRi knockdown (vulnerability index −5.7, 95% CI −6.2 to −5.2) (Bosch et al., 2021), detected by mass spectrometry, near-invariant across clinical isolates (only five polymorphic sites over 145,209 genomes, none disruptive), and predicted to be very well folded (mean pLDDT 95.3). This invariance holds at deeper phylogenetic scales: Rv1025 differs from the immediate outgroup *M. canettii* by a single, synonymous substitution, and it is a genus-core gene retained across every sampled *Mycobacterium* species. (Given only five intra-complex sites and one outgroup substitution, we report these as counts of segregating/substituted sites rather than as *dN/dS* or *pN/pS* ratios, which are underpowered at this sampling and would over-interpret a one-variant swing.) Yet neither Rv1025 nor its family has a functional annotation. The Pfam family DUF501 (PF04417, InterPro IPR007511) is described only as “family of uncharacterised bacterial proteins”, comprising 4,370 proteins across 5,484 taxa with no associated Gene Ontology term and, critically, no experimentally solved structure; the eggNOG orthology (COG1507) and KEGG orthology (K09009) are likewise “function unknown”. The essentiality, however, is not idiosyncratic to *Mycobacterium tuberculosis*: the DUF501 gene is reported essential in three Actinobacterial genera, *Mycobacterium tuberculosis* (Rv1025), *Corynebacterium glutamicum* (cgp_1113) and *C. diphtheriae* (DIP0919) (Goodall et al., 2023), identifying DUF501 as a conserved, non-redundant core function whose molecular nature is unknown.

### 3.2 DUF501 has no detectable structural or sequence-profile homolog

To seek a function from structure, we searched the high-confidence AlphaFold model of Rv1025 against complete structural databases with Foldseek. No significant homolog was found: the AlphaFold/Swiss-Prot search returned no hit, PDB100 returned a single partial and non-significant match (26% of the query at 24% identity), and CATH50 returned five marginal hits (13–19% identity, *e* ≥ 0.01) scattered over five *different* folds without convergence. A complementary profile–profile search with HHpred, which is more sensitive to remote homology, likewise returned no significant hit beyond the family itself: the only high-probability matches were self-hits to the DUF501 profiles (COG1507 and PF04417, probability 100%), while every other hit fell below 56% probability with E-values ≥ 18 and did not converge on any characterised fold. DUF501 therefore adopts a fold that is not represented in the PDB or in CATH and has no detectable sequence-profile relative; the Rv1025 model is, to our knowledge, the first structural representative of the family. Homology-based transfer cannot assign its function.

### 3.3 The *eno*–*divIC* –*Rv1025* –*ppx2* operon is conserved across the Actinobacteria phylum

Although the operon mixes unrelated pathways, its arrangement is nonetheless strongly conserved. Querying the four proteins against 63 reference genomes, the complete four-gene block was found intact and in the same order, with *divIC* always the immediate neighbour of *Rv1025*, in 61 of 63 genomes (Figure 1): the entire genus *Mycobacterium* (48/48 species, including the reductive genomes of *M. leprae* and *M. lepromatosis*, which retain the whole block), the order Corynebacteriales (*Corynebacterium, Nocardia, Rhodococcus, Gordonia, Tsukamurella, Mycobacteroides*), and representatives of Streptomycetales, Bifidobacteriales, Propionibacteriales and Micrococcales. DUF501 was entirely absent from the two out-of-phylum negatives (*E. coli, B. subtilis*). Consistently, the family is strongly enriched in this phylum without being confined to it: of the 4,378 DUF501 proteins currently in InterPro, 3,830 (87%) are Actinomycetota, the remainder being a minority of Pseudomonadota (264), scattered eukaryotic entries (179) and a handful of Bacillota (24) (Paysan-Lafosse et al., 2023). Conservation of an ordered gene block over hundreds of millions of years, in genomes whose gene order is otherwise extensively rearranged, points to a functional association among these genes rather than passive co-inheritance. To check that this is not merely the genome-wide baseline of gene-order conservation, we applied the identical synteny test to 90 random four-gene H37Rv blocks across the eleven distant, non-*Mycobacterium* Actinobacteria, conditioning on orthology so that only arrangement, not gene conservation, is scored (Methods). Among the 43 blocks whose genes are broadly conserved, synteny given orthology is preserved at a median of 0.55 (mean 0.47), against 1.00 for the operon: the *eno*–*divIC* –*Rv1025* –*ppx2* block is fully syntenic wherever its genes are present and sits in the top fifth of conserved blocks (8 of 43 as syntenic or more; Supplementary Table S4). Its arrangement is therefore maintained well above the passive-co-inheritance baseline, although, as expected for the operon-rich Actinobacteria, it is not a unique outlier. The block is retained from *Mycobacterium tuberculosis* to *Bifidobacterium* (Rv1025 identity 100% to 51%) along the Actinobacterial taxonomy (Supplementary Figure S3).

**Figure 1:**
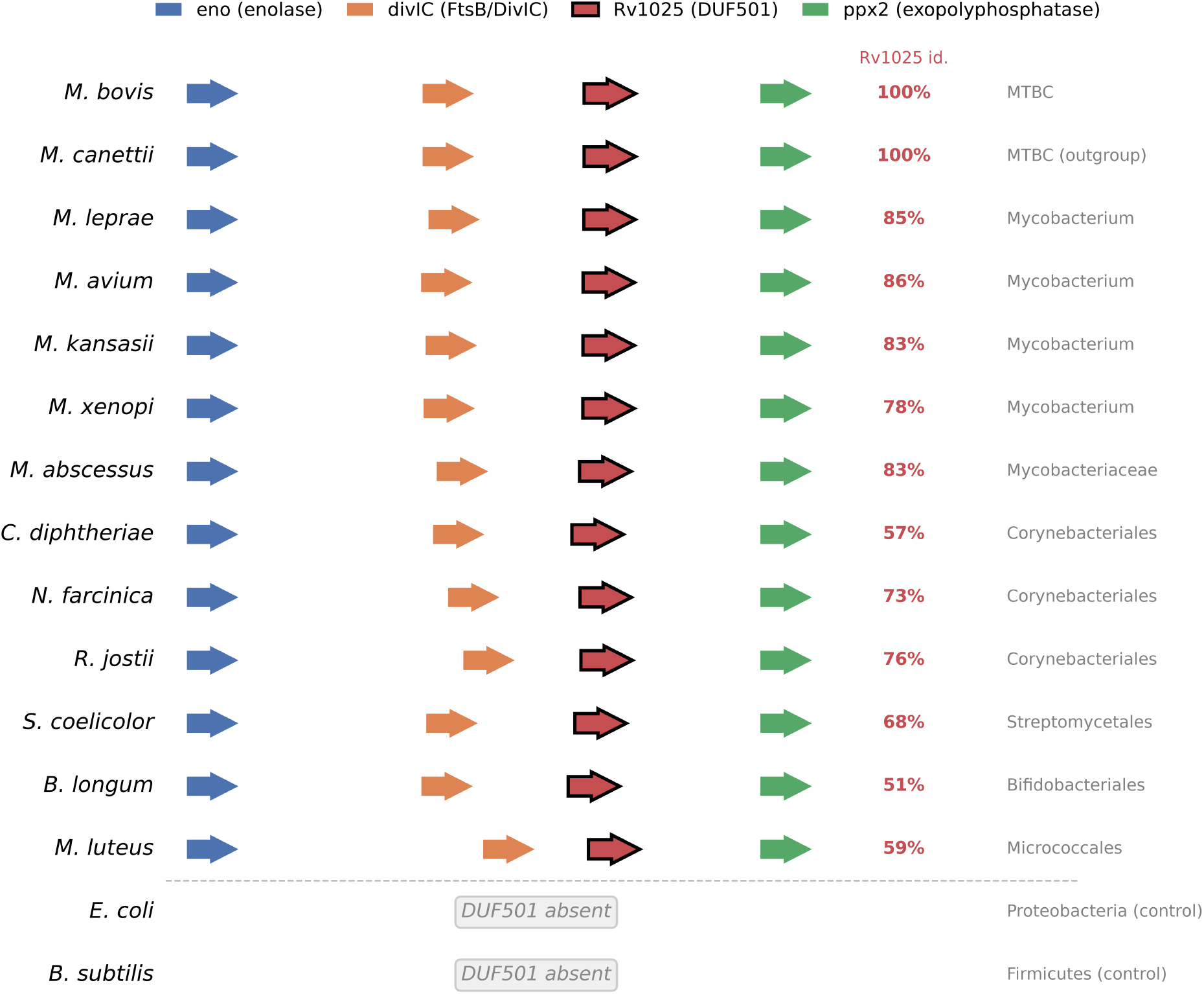
The *eno*–*divIC* –*Rv1025* –*ppx2* operon is a conserved genomic unit across the Actinobacteria. Representative genomes are ordered from the *M. tuberculosis* complex (top) to distant Actinobacterial orders, with two out-of-phylum negatives at the bottom. Gene order is normalised so the operon reads left to right (in several genomes, e.g. *N. farcinica, S. coelicolor*, the block lies on the opposite strand). *divIC* (FtsB/DivIC family) is the invariant immediate neighbour of *Rv1025* (DUF501, outlined). The Rv1025 amino-acid identity to *Mycobacterium tuberculosis* is shown at right; DUF501 is absent from *E. coli* and *B. subtilis*. Full data for all 63 genomes are in Supplementary Table S1.

### 3.4 Rv1025 does not form a stable binary complex with DivIC

Because DivIC (FtsB/DivIC family) is the invariant syntenic neighbour, we tested a direct physical interaction with AlphaFold-Multimer. The prediction was negative and specific. As a positive control, the established divisome pair DivIC–FtsQ was recovered reproducibly (ipTM 0.38 across the five models, minimum inter-chain predicted aligned error 5.1 Å), showing that the assay can detect a genuine division interface, albeit with a modest ipTM ceiling typical of small membrane-protein interfaces. Against this calibration, Rv1025–DivIC failed: its best ipTM was 0.25 but collapsed to 0.08 across models (no converged pose), with an inter-chain error of 14.7 Å, at the level of the specificity controls Rv1025–Enolase (0.19) and Rv1025–Ppx2 (0.17). The deep syntenic conservation therefore reflects a functional association or co-regulation, not a demonstrable stable heterodimer with DivIC (Figure 2).

**Figure 2:**
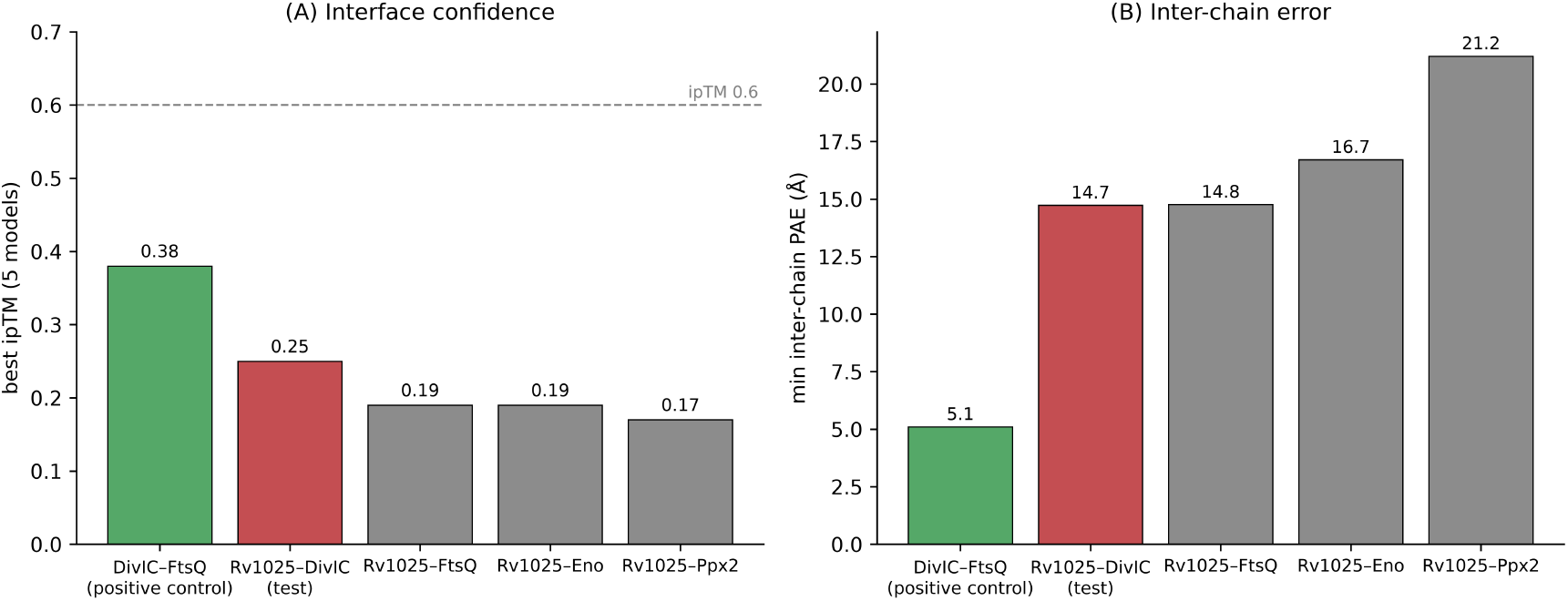
AlphaFold-Multimer does not support a direct Rv1025–DivIC complex. **(A)** Best interface pTM (ipTM) over the five models and **(B)** minimum inter-chain predicted aligned error (PAE) for each pair. The established divisome pair DivIC–FtsQ (positive control, green) is the only one combining a higher ipTM with a low inter-chain PAE (5.1 Å), indicating a genuine interface; the Rv1025–DivIC test (red) and the specificity controls (Rv1025 with FtsQ, Enolase, Ppx2; grey) all show low ipTM and high inter-chain PAE.

### 3.5 A near-invariant Cys–His–Glu triad forms a metal site supported by holo AlphaFold3

We then turned to conservation. Over an alignment of 8,700 homologous sequences, 23 residues are near-invariant (≥ 90% identity to the query); among them, several of catalytic type, notably Cys113 and His115 (both 100% conserved) and Glu59 (99%). The invariance of the three residues also holds on the curated Pfam seed alignment (39 sequences), independent of the redundancy of the large alignment, where Cys113, His115 and Glu59 are each fully conserved (Supplementary Figure S1). On the AlphaFold model these three residues form a compact cluster within a cleft. Measuring their geometry, the side-chain donor atoms (Cys-S*γ*, His-N*δ*1, Glu-O*ε*) converge on a common point ~2.3 Å from each, the hallmark of a mononuclear metal-coordination site rather than a linear catalytic-triad relay.

To test this, we ran holo AlphaFold3 predictions of Rv1025 with a single Zn, Fe or Mn ion. In every case the ion was placed in the Cys113–His115–Glu59 site at canonical metal–ligand distances and with very high per-ion confidence (Table 1; Figure 3): 2.25–2.47 Å to the three donors, per-ion pLDDT 94–98, reproducibly, with iron giving the most confident model. The protein coordination is incomplete: only three protein ligands (with a possible bidentate contribution of Glu59) engage the metal, leaving open coordination positions facing the pocket lined by the conserved, charged second-shell residues Lys112 and Arg92. Such an incompletely protein-coordinated, solvent-open metal is more consistent with a catalytic metal centre than with a purely structural one. As a specificity control, we mutated the three ligands to alanine (C113A/H115A/E59A) and repeated the holo prediction. The fold was preserved (mean pLDDT 96), but the metal no longer occupied the site: it relocated 16–22 Å away to a non-conserved surface cluster (Cys20/His48/Glu24, at 74/20/8% family conservation versus 99–100% for the true triad). Abolishing the conserved triad therefore abolishes metal binding at that site, indicating that the wild-type placement is specific to Cys113–His115–Glu59 rather than an arbitrary surface artefact. The relocation also shows that AlphaFold3 will accommodate a divalent metal at any suitable Cys/His/Glu cluster, so per-ion confidence alone is not evidence of physiological occupancy, which will require experimental metallation.

**Table 1:** Holo AlphaFold3 coordination of the Cys113–His115–Glu59 site by three divalent metals (top-ranked model). Distances in Å; pLDDT is the per-ion confidence.

| Metal | per-ion pLDDT | Cys113 Sγ | His115 Nδ1 | Glu59 Oε | triad ligands |
| --- | --- | --- | --- | --- | --- |
| Fe | 98.3 | 2.37 | 2.31 | 2.27 | 3/3 |
| Mn | 97.6 | 2.47 | 2.38 | 2.29 | 3/3 |
| Zn | 94.6 | 2.40 | 2.32 | 2.25 | 3/3 |

**Figure 3:**
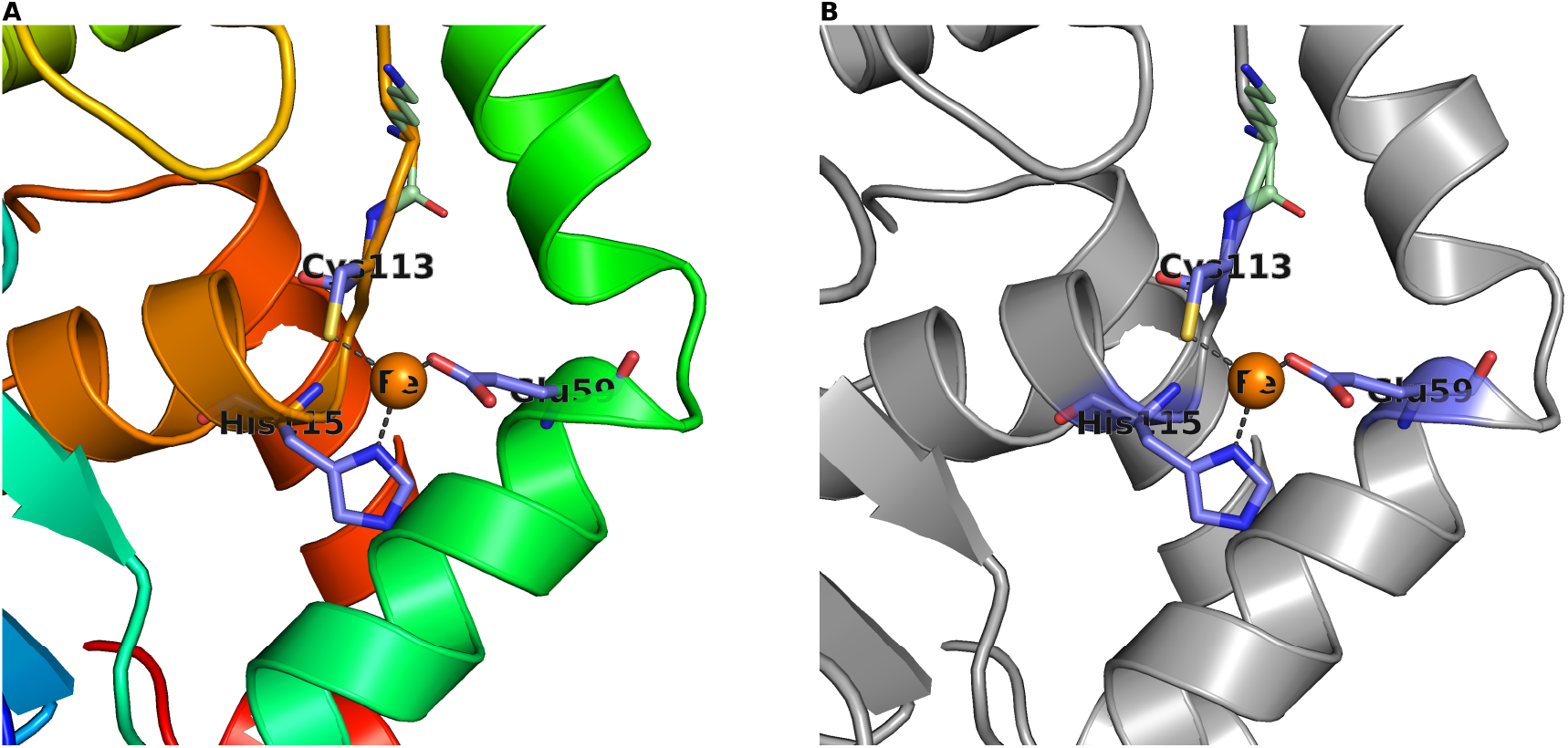
The predicted metal site of Rv1025 (DUF501). **(A)** Overall AlphaFold fold (rainbow, N to C) with the three metal-coordinating residues shown as sticks; the family has no experimentally solved structure. **(B)** Close-up of the holo AlphaFold3 model with iron: the metal (orange sphere) is coordinated by the near-invariant Cys113, His115 and Glu59 (dashed bonds), with open positions facing the conserved second-shell pocket. Rendered with PyMOL.

We sought orthogonal support that does not rely on AlphaFold’s learned metal placement. BioMetAll, which scores metal-binding sites from backbone preorganisation alone, was run blind on the apo model and independently returned the Cys113–His115–Glu59 triad as a metal-compatible site, with a predicted centre at coordination distance from the donor atoms. Consistent with the mutation experiment, it also returned the non-conserved Cys20/His48/Glu24 cluster: geometry alone therefore identifies both cavities, and it is the deep conservation of the Cys113–His115–Glu59 triad (99–100% versus 8–74% for the surface cluster), not its geometry, that singles it out as biologically meaningful. Two orthogonal predictors thus converge on the same site while agreeing that backbone geometry is permissive at more than one location (Supplementary Table S2). A template-based search with AlphaFill returned no holo homolog from which to transplant an ion, as expected for a family with no structural relative, so it provides no independent metal evidence either way.

Pocket detection on the apo model with fpocket found that the conserved metal site forms a distinct, well-defined cavity (255 Å^3^, 22 alpha spheres; alpha-sphere centres within 4.4 Å of the triad donors) and that the fold additionally presents a separate, clearly druggable pocket (druggability score 0.65, 361 Å^3^), so the protein is structurally tractable rather than flat or featureless (Supplementary Figure S2, Supplementary Table S3). That second cavity is not an incidental dent of this particular model: the fourteen residues lining it are conserved at 74% across the family alignment, against a 54% whole-protein background and above 99.5% of size-matched random residue sets (*p* = 0.005), approaching the conservation of the metal-site pocket itself (78%). The contrast is not an artefact of alignment redundancy: applying Henikoff position-based sequence weights (Henikoff and Henikoff, 1994), which reduce the alignment to 2,599 effective sequences out of 8,700, leaves the pocket at 67% against a 51% background (*p* = 0.011). A conserved, druggable cavity distinct from the metal centre is therefore a second candidate functional site, and a second potential point of intervention. Mapping all 23 near-invariant residues onto the model supports the same reading while keeping it modest: twenty of them form a single buried cluster (mean relative solvent accessibility 0.18) that comprises the metal triad and is otherwise ordinary fold-core constraint, whereas the only solvent-exposed conserved group (mean 0.49) is a three-residue N-terminal patch, two of whose members line this second pocket. That patch is itself near-invariant (92% after redundancy weighting) and, like six of the fourteen pocket residues, falls outside the annotated PF04417 domain, which spans residues 17–136: the second cavity is thus built partly from a conserved N-terminal extension that the domain definition does not cover. Because that segment is also the least confident part of the model (per-residue pLDDT 81–87, against 95.3 overall), we treat it as consistent with, rather than evidence for, a functional role. The metal cavity itself received only a modest fpocket druggability score (0.24), the expected behaviour of a hydrophobicity-weighted scoring function on a polar metal site rather than evidence against druggability. Geometric measurement of coordination-sphere accessibility, which does not depend on that scoring function, instead argues positively for it. Excluding water and any other non-protein atom, Rv1025 opens 19.8% of its coordination sphere to a probe at the 2.4 Å threshold, 14.3% of it facing the solvent – against 10.1–13.9% open and 0.5–4.9% solvent-facing for three validated catalytic zinc targets (carbonic anhydrase II, thermolysin, LpxC) and a clean negative control at 0.0%/0.0% for the structural, non-catalytic zinc finger Zif268, which validates the measurement itself (Methods). The Rv1025 metal site is therefore at least as chemically accessible to an exogenous ligand as established drug targets, and nothing like the saturated, buried geometry of a structural zinc. This is a property of the fold, not of one model: accessibility is essentially unchanged across ion identity (Zn 17.2%, Fe 19.8%, Mn 20.6%) and across the five AlphaFold3 models per ion (19.8% ± 0.3% for Fe). A possible confound – AlphaFold models being more loosely packed than crystal structures, which would inflate open space regardless of any true difference – was addressed directly by repeating the measurement on AlphaFold DB models of the two calibration proteins with a crystallographic active site: the AlphaFold-versus-crystal shift is small and inconsistent in sign (carbonic anhydrase II 7.4% versus 12.2%; thermolysin 16.2% versus 13.9%), too weak and too erratic to explain the much larger and unidirectional gap seen for Rv1025 (*n* = 2; treated as a weak, non-systematic bias rather than a negligible one). This establishes chemical accessibility of the site, not affinity, selectivity, or that greater openness is pharmacologically better – a more exposed metal buries less surface and could bind a small chelator more weakly.

Because conservation alone cannot separate a functional constraint from ordinary fold constraint, we asked whether residue coevolution could resolve the functional site any further. Direct coupling analysis of the same alignment was well powered (2,260 effective sequences for 155 positions) and behaved as it should: 58% of the top *L/*2 predicted pairs are genuine contacts in the model, against a 2.8% background rate, a 20.6-fold enrichment. The signal, however, is almost entirely the fold itself. Among the 155 strongest couplings, only 16 join residues more than 15 Å apart in the monomer, where chance alone would place 112 (*p* = 6 × 10^−59^), so coevolution adds no functional map beyond the triad. This reflects the protein rather than the method: the three metal ligands are 99% invariant, and an approach that measures covariation is structurally blind to positions that do not vary.

Two specific questions could nonetheless be settled. First, the conservation of the druggable pocket is not a spillover from the adjacent metal site: across 80 eligible pocket-to-pocket pairs, the mean coupling (−0.013) does not exceed a distance-matched null (−0.038; *p* = 0.23), so the two cavities are not evolutionarily coupled and the conservation of the second pocket is an independent constraint. Second, the incomplete protein coordination of the metal raises the possibility that a second subunit completes the site in trans; a conserved homodimer interface would leave strong couplings unexplained by the monomer, carried by exposed residues grouped into a surface patch. Neither expectation is met: the 25 residues carrying those couplings are no more exposed than the rest of the protein (mean relative solvent accessibility 0.24 against 0.27; *p* = 0.84) and form a diffuse set spanning residues 5 to 124 rather than a patch. We therefore find no evidence for a conserved oligomeric interface, while noting that a recent or non-obligate oligomer would leave no coevolutionary signature, so the oligomeric state remains a question for experiment (Supplementary Table S8).

### 3.6 The metal site is a universal signature of bacterial DUF501

The evidence above establishes a metal site in one protein. Whether that site defines the family, or is a mycobacterial specialisation within it, is a different question, and one that decides how far the functional hypothesis can be carried. We therefore examined the full Pfam alignment of PF04417 (2,228 sequences, 471 columns), in which Rv1025 is present as P96375/16–137, so that the three positions can be located by direct anchoring rather than by realignment.

The triad is strictly universal in bacteria. Every one of the 1,472 near-complete bacterial sequences carries Glu, Cys and His at the three positions (100.0%), and the alignment as a whole contains no non-conservative substitution at any of them: not a single Glu-to-Asp, and no replacement of either Cys113 or His115 (Supplementary Table S9). The 86 apparent exceptions are gaps rather than substitutions, and 75 of them fall in sequence fragments (median aligned length 76 residues against 124 for the rest, *p* = 1 × 10^−46^). The metal site is therefore not a mycobacterial peculiarity but the defining feature of the family in bacteria, which is what allows a functional hypothesis derived from Rv1025 to be stated for DUF501 as a whole.

The family also extends beyond bacteria. It includes 102 near-complete eukaryotic members, essentially protists, among them apicomplexan parasites (*Babesia, Neospora, Eimeria*), stramenopiles and *Naegleria*, in which the domain is frequently embedded within a larger protein. These are genuine homologs rather than marginal profile matches, at a mean 30% identity to Rv1025 over the domain. Eleven of them have lost part of the site, almost always the Cys–His pair with retention of the glutamate (*p* = 5 × 10^−14^ against bacteria), and this is not explained by divergence, since those eleven are no less similar to Rv1025 than the ninety-one that retain the complete triad (*p* = 0.36). We report this eukaryotic branch as an observation of sequence and do not interpret it further here: whether these proteins bind a metal by other means, or not at all, is untested, and the proportion affected is an upper bound because 576 sequences of the alignment could not be assigned a lineage (all of which carry the complete triad).

## 4 Discussion

Bringing these results together, we propose that DUF501 is a metal-binding protein and candidate metalloenzyme. The evidence is convergent: a fold with no structural relative (ruling out annotation by homology), a strictly conserved and essential gene across Actinobacterial genera, and a near-invariant Cys–His–Glu constellation whose apo geometry and holo behaviour both indicate a divalent-metal centre, plausibly catalytic. Catalysis is one hypothesis among others (metal storage, transfer or sensing, or a structural or redox role), to be distinguished experimentally. The presence of a cysteine thiol ligand is informative: thiolate coordination points to a transition-metal, possibly redox-active site, and is consistent with the weak, non-significant structural signals previously observed for Rv1025 towards metal- and iron-handling proteins. To our knowledge this is the first functional hypothesis proposed for the DUF501 family, and it illustrates a general route to illuminating a dark protein family through a single tractable, well-folded member when structural homology fails: population-scale conservation mapped onto a high-confidence model, adjudicated by holo modelling.

Several questions remain open. AlphaFold3 accommodates all three metals and does not, by itself, identify the physiological one, although iron yields the most confident model and the thiolate ligand is compatible with iron. The precise reaction catalysed is not established; the pocket architecture, with a conserved charged second shell (Lys112, Arg92), is compatible with a hydrolytic or redox activity acting on a small charged substrate, but this requires biochemical testing. The conserved operon association with a glycolytic enzyme, a division protein and an exopolyphosphatase does not point to a shared pathway: each protein’s strongest predicted partners lie in its own process (enolase with pyruvate kinase, DivIC with the divisome protein FtsQ), no transcription-factor regulon is shared across the four genes in the *Mycobacterium tuberculosis* regulatory network, and the only within-block association, Rv1025 with Ppx2, is carried by genomic-context (neighbourhood and gene-fusion) channels rather than by experimental evidence. The block is thus co-transcribed but functionally heterogeneous, and the conservation of its arrangement reflects operon-level selection rather than a common biochemical function; our negative complex prediction argues specifically against a stable Rv1025–DivIC heterodimer, without excluding a transient or divisome-context interaction. A wider Boltz-2 screen against five further divisome components (FtsZ, FtsW, PbpB/FtsI, FtsK, SepF) was uniformly negative once judged on inter-chain PAE rather than raw ipTM alone: the highest nominal ipTM in the panel (FtsW, 0.281) exceeded the in-panel positive control (DivIC–FtsQ, 0.221), yet its inter-chain PAE (18.4 Å) was close to double the control’s (10.4 Å), the same ipTM-without-PAE trap that the calibration protocol was designed to catch. Rv1025 therefore shows no evidence of a stable binary complex with any divisome component tested. Separately, a direct AlphaFold3 test of a Rv1025 homodimer, with and without the two holo metal ions, likewise gave no confident interface (ipTM 0.16, at the level of the monomer specificity controls), converging with the coevolutionary analysis above on the absence of an obligate oligomeric interface able to complete the metal coordination sphere in trans; the oligomeric state therefore remains an open, experimentally tractable question rather than a structural inference.

From a translational standpoint, Rv1025 unites the features sought in an antitubercular target: essentiality confirmed across genera, strong CRISPRi vulnerability, near-invariance across clinical isolates (limiting pre-existing resistance), an ordered fold that presents a druggable pocket, and a defined, conserved metal cavity that is geometrically as accessible to an exogenous ligand as established zinc-dependent drug targets. Metal sites are classically amenable to inhibition, including by metal-chelating warheads, and the polyphosphate pathway to which the operon’s exopolyphosphatase Ppx2 belongs has already yielded a validated antitubercular target in the polyphosphate kinase PPK2 (Singh et al., 2016). DUF501 therefore offers a double return: a candidate drug target in *Mycobacterium tuberculosis* and the functional entry point to an entire, previously dark bacterial protein family.

## 5 Conclusion

Rv1025 exemplifies an essential, vulnerable, structurally ordered but functionally dark *Mycobacterium tuberculosis* protein. We show that its family, DUF501, has a novel fold and a deeply conserved operon context, that it does not form a stable binary complex with its syntenic division neighbour, and that it carries a near-invariant Cys113–His115–Glu59 metal site supported by holo AlphaFold3. We propose that DUF501 is a metal-binding protein, plausibly a metalloenzyme, and a candidate antitubercular target. These conclusions are computational; experimental determination of the bound metal, the structure and the catalysed reaction is the natural next step, and would characterise, in one move, a protein family conserved across the Actinobacteria.

## Supporting information

Supplementary Figure S1: Pfam seed alignment conservation of the metal triad

Supplementary Figure S2: Pocket detection and druggability

Supplementary Figure S3: Synteny across the Actinobacteria phylum

Supplementary Table S1: Synteny across 63 reference genomes

Supplementary Table S2: Orthogonal metal-site prediction (BioMetAll)

Supplementary Table S3: Pocket druggability scores

Supplementary Table S4: Synteny null model (90 random blocks)

Supplementary Table S5: Pocket residue conservation

Supplementary Table S6: Near-invariant residue clusters

Supplementary Table S7: Henikoff-weighted conservation

Supplementary Table S8: Pocket coupling analysis

Supplementary Table S9: DUF501 family triad universality

Supplementary Table S10: DUF501 in eukaryotic sequences

## Declarations

### Funding

This research did not receive any specific grant from funding agencies in the public, commercial, or not-for-profit sectors.

### Competing interests

The author declares no competing interests.

### Ethics approval

Not applicable. This study did not involve human participants, human data or tissue, or animals.

### Consent to participate

Not applicable.

### Consent for publication

Not applicable.

### Availability of data and materials

All data analysed in this study are publicly available and are cited at each use throughout the Materials and Methods, with accession numbers and database sources given in place. The principal sources are the H37Rv reference annotation (NC_000962.3), the AlphaFold model of Rv1025 (AlphaFold Protein Structure Database, AF-P96375-F1), the InterPro/Pfam family PF04417 and its full alignment, UniProt, and the published transposon and CRISPRi essentiality screens cited in the text.

### Code availability

Analysis scripts (synteny, conservation, AlphaFold and Boltz-2 job generation and parsing, figure rendering) and intermediate results are deposited at https://github.com/cguyeux/rv1025-duf501-mtbc.

### Authors’ contributions

Christophe Guyeux: Conceptualization, Data curation, Formal analysis, Investigation, Methodology, Project administration, Resources, Software, Validation, Visualization, Writing: original draft, Writing: review and editing.

### Use of generative AI and AI-assisted technologies in the writing process

During the preparation of this work, the author used Claude Code (Anthropic), an AI-agent software environment powered by the Claude Sonnet 5 model, for in silico data retrieval, statistical analysis, literature verification and drafting assistance, as described in the Materials and Methods. After using this tool, the author reviewed, independently verified against primary sources, and edited the content as necessary, and takes full responsibility for the content of this publication.

## Notes

https://github.com/cguyeux/rv1025-duf501-mtbc

### Competing Interest Statement

The authors have declared no competing interest.

