## Supplementary figures and images for "A conserved cysteine–histidine–glutamate metal site identifies DUF501 (Rv1025), an essential uncharacterised protein family of *Mycobacterium tuberculosis*, as a candidate metalloenzyme and drug target"

### Supplementary Figure S1: Pfam seed alignment conservation of the metal triad

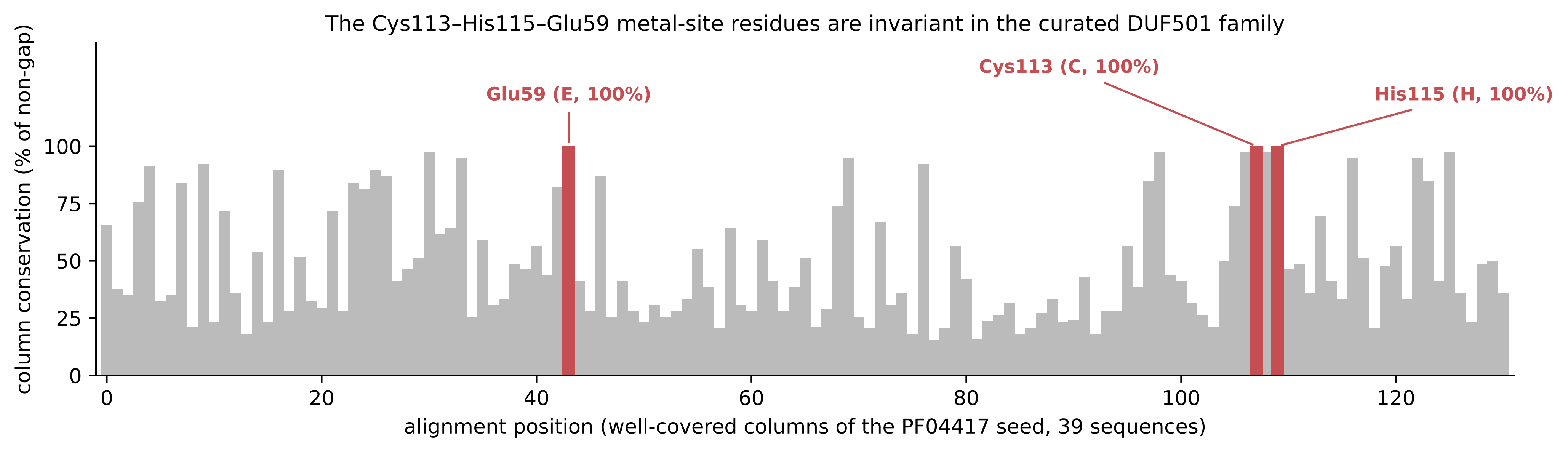

### Supplementary Figure S2: Pocket detection and druggability

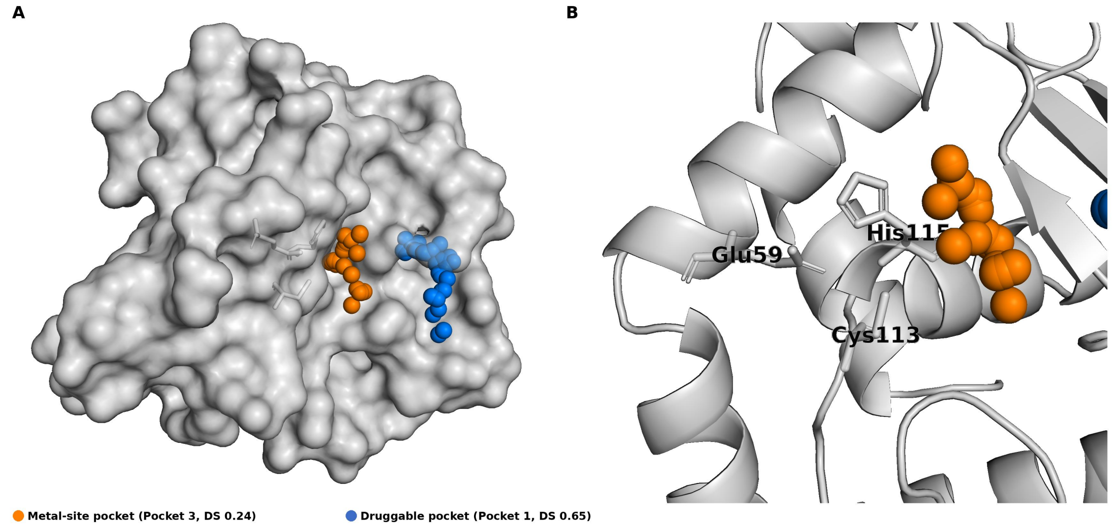

### Supplementary Figure S3: Synteny across the Actinobacteria phylum

The eno–divIC–Rv1025–ppx2 block is retained across the Actinobacteria phylum

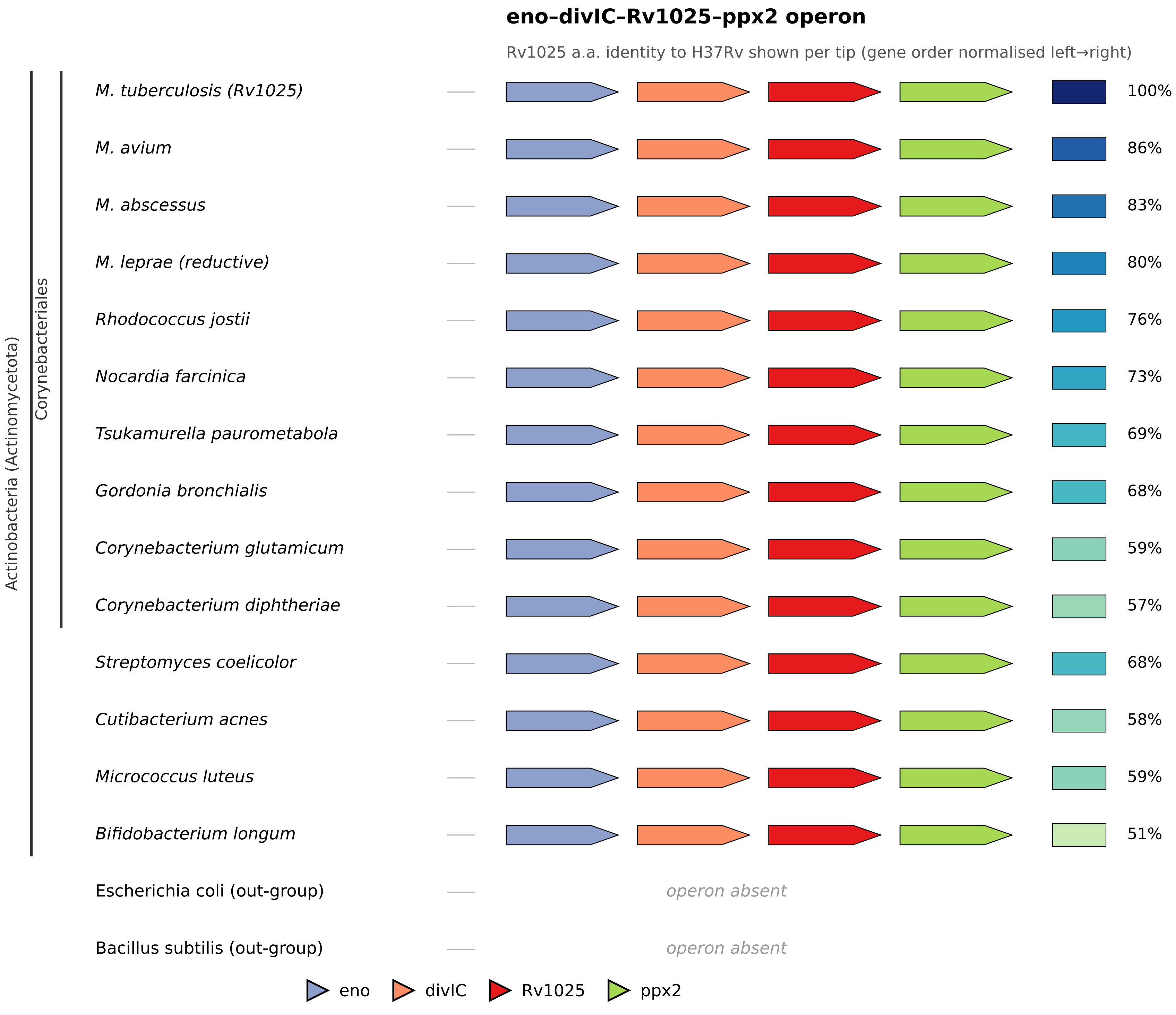
